# Host-microbiome mutualism drives urea carbon salvage and acetogenesis during hibernation

**DOI:** 10.1101/2025.02.13.638127

**Authors:** Matthew D. Regan, Edna Chiang, Michael Grahn, Marco Tonelli, Fariba M. Assadi-Porter, Garret Suen, Hannah V. Carey

## Abstract

2.

Hibernation is a seasonal survival strategy employed by certain mammals that, through torpor use, reduces overall energy expenditure and permits long-term fasting. Although fasting solves the challenge of winter food scarcity, it also removes dietary carbon, a critical biomolecular building block. Here, we demonstrate a process of urea carbon salvage (UCS) in hibernating 13-lined ground squirrels, whereby urea carbon is reclaimed through gut microbial ureolysis and used in reductive acetogenesis to produce acetate, a short-chain fatty acid (SCFA) of major value to the host and its gut microbiota. We find that urea carbon incorporation into acetate is more efficient during hibernation than the summer active season, and that while both host and gut microbes oxidize acetate for energy supply throughout the year, the host’s ability to absorb and oxidize acetate is highest during hibernation. Metagenomic analysis of the gut microbiome indicates that genes involved in the degradation of gut mucins, an abundant endogenous nutrient, are retained during hibernation. The hydrogen disposal associated with reductive acetogenesis from urea carbon helps facilitate this mucin degradation by providing a luminal environment that sustains fermentation, thereby generating SCFAs and other metabolites usable by both the host and its gut microbes. Our findings introduce UCS as a mechanism that enables hibernating squirrels and their gut microbes to exploit two key endogenous nutrient sources – urea and mucins – in the resource-limited hibernation season.

**SIGNIFICANCE STATEMENT:** When food becomes scarce during winter, hibernating mammals induce torpor to minimize energy demands and enable monthslong fasting. However, fasting eliminates the intake of essential nutrients such as carbon. We identified a two-step microbial-host interaction in ground squirrels – urea carbon salvage (UCS) – which counters carbon limitation by salvaging carbon from waste urea. Through activities of ureolytic and acetogenic bacteria, urea-derived CO_2_ is reduced by free hydrogen to form acetate, whose oxidation provides energy for gut microbes and the host. This process also helps maintain a permissive environment for fermentation of other host-derived, energy-dense compounds such as mucins. UCS broadens our understanding of host-microbe mutualism under extreme nutritional constraints and may represent a widespread adaptation among fasting mammals.

## Introduction

For most mammals and their gut microbes, winter is a time of resource limitation. Some species survive by storing extra fat reserves and finding other, less preferred food items. In contrast, hibernators like the 13-lined ground squirrel (*Ictidomys tridecemlineatus*) survive by entering torpor for weeks at a time, which allows them to stop eating for approximately six months each year [1]. Winter fasting solves the hibernator’s food scarcity problem, but it deprives the animal – and its gut microbes – of exogenous nutrients that are crucial for sustaining life. This challenge could be met in part by mutualistic relationships between hosts and their microbial symbionts that transform endogenous compounds into valuable nutrient resources.

One example of resource reclamation during fasting is urea nitrogen salvage (UNS), a gut microbe-dependent process used by ground squirrels to recycle urea nitrogen back into their tissue protein pools during hibernation [2], when they lack the typical dietary supply of nitrogen. This process, which peaks during late winter after months of fasting and just prior to the squirrel’s emergence into the spring mating season, recycles nitrogen to support tissue protein synthesis throughout winter, and likely contributes to these hibernators’ renowned resistance to muscle atrophy [2,3]. Urea nitrogen is also incorporated into the microbial protein pool, thereby confirming that UNS benefits both symbiotic partners [2]. Thus, urea, which is excreted as a waste product during summer, serves as an endogenous nitrogen source during hibernation when these animals and their gut microbes lack an exogenous supply of this important biomolecular building block.

In addition to nitrogen, urea also contains carbon, another essential nutrient that is limited during hibernation due to long-term fasting. Theoretically, the ability to capture this carbon via urea carbon salvage (UCS) would also provide nutritional benefits to the microbiota and potentially to the hibernator host. Although various animals are known to use UNS as a nitrogen conservation system [4], to our knowledge UCS has never been proposed as a mechanism to maintain carbon balance in species that undergo extended fasting, which is the case for most hibernators. But preliminary results from our previous study [2] support the existence of UCS and suggest that urea carbon is liberated by microbial urea hydrolysis, biologically fixed, and incorporated into organic molecules. Thus, it is possible that both host and microbes benefit from UCS during resource-limited hibernation.

In this study, we tested more rigorously the hypothesis that carbon liberated by urea hydrolysis is incorporated into organic molecules, and that these molecules provide benefits for ground squirrels and/or their microbiota during winter fasting. To accomplish this, we treated squirrels with ^13^C,^15^N-labeled urea and measured urea carbon (i.e., ^13^C) incorporation into the microbiota as well as tissue metabolome and protein. We performed real-time in vivo δ^13^C breath experiments to estimate rates of microbial urea hydrolysis and absorption/metabolism of orally gavaged acetate, as well as metagenomic analyses of bacterial genes involved in carbon fixation and acetogenesis. In addition, we expanded the notion that fasted hibernators and their gut microbes participate in mutualistic interactions that enable the use of host-secreted compounds as nutrients by examining whether hibernator microbiomes are enriched in genes encoding enzymes that degrade intestinal mucins, which are the most abundant endogenous nutrients in the squirrel gut ecosystem during winter fasting.

## Materials & methods

### Experimental animals

From a total of 65 13-lined ground squirrels (39 females, 26 males; hereafter called squirrels), 39 were used for the doubly labelled urea treatment experiments described in [2], 15 were pretreated with oral gavages of ^13^C-acetate, and gut contents from 11 others were used for carbon fixation and mucin degradation metagenomics analyses. All squirrels were born in a vivarium on the University of Wisconsin-Madison (UW-Madison) campus to wild-caught female squirrels that were collected around Madison, Wisconsin, USA in May 2018 and May 2019. The pregnant squirrels were housed individually at 22°C with a 12:12 h light-dark cycle with ad libitum access to water and rodent chow (Teklad no. 2020X, Envigo, Indianapolis, IN, USA) supplemented with fruit (apples) and sunflower seeds once per week. After birth, pups remained with their mothers for 5 weeks and then moved to individual cages. Following 2 weeks of ad libitum chow and fruit, food was restricted to 12 g chow per day (supplemented with 1 g of sunflower seeds once per week) to prevent excessive weight gain that occurs in captive-born squirrel pups. The consistent diet minimized diet-induced effects on the gut microbiota and the breath δ^13^C value. Squirrels were held under these conditions until the day of experiment (Summer group) or transferred to the cold room at the beginning of the hibernation season (Early and Late Winter groups).

In mid-September, squirrels in the Winter groups were moved to a 4°C environmental chamber in the UW-Madison Biotron. The chamber was held in constant darkness except for ∼5 min/day of dim light to enable activity checks. Food and water were removed after squirrels began regular torpor-arousal cycles, which was 24–72 h after being moved into the chamber.

Each of the 39 urea-treatment squirrels were studied at one of three time points: Summer (14 squirrels), Early Winter (12 squirrels), and Late Winter (13 squirrels). Each seasonal group was divided into three treatment groups: ^13^C,^15^N-urea with antibiotics, ^13^C,^15^N-urea without antibiotics, and unlabeled urea without antibiotics. Mean days in hibernation for the Early Winter and Late Winter groups were 36 d (range 31–41 d) and 127 d (range 109–135 d), respectively. Mean body masses on experiment days for Summer, Early Winter, and Late Winter groups were 187.1 g (range 165.0–224.0 g), 155.6 g (range 123.4–187.2 g), and 134.1 g (range 105.5–167.0 g), respectively.

The 15 acetate-treatment squirrels were divided into four groups: Summer with antibiotics (4 squirrels), Summer without antibiotics (3 squirrels), Late Winter with antibiotics (3 squirrels), and Late Winter without antibiotics (5 squirrels). Mean days in hibernation for the combined Late Winter groups was 92 d (range 68–145 d). Mean body masses on experiment days for Summer and Late Winter were 206.1 g (range 172.6–241.2 g) and 151.2 g (range 128.3–174.0 g), respectively.

The 11 microbiota-intact squirrels used for metagenomic analysis of carbon fixation and mucin degradation genes included 6 Summer and 5 Late Winter squirrels. Mean days in hibernation for the Late Winter group were 135 d (range 127–144 d). Mean body masses on experiment days for Summer and Late Winter were 183.0 g (range 144.0–225.0 g) and 132.5 g (range 116.5–144.6 g) The UW-Madison Institutional Animal Care and Use Committee approved all procedures.

### Animal treatments and stable isotope breath-testing

For the 39 squirrels treated with ^13^C,^15^N-labeled urea, the antibiotic treatments to deplete the gut microbiota, the ^13^C,^15^N-labeled urea treatments (200 mg urea kg^-1^ body mass), the breath testing experiments via cavity ring-down spectrometry (CRDS), and the euthanasia protocol were all carried out as described in [2].

The 15 squirrels used to estimate host and microbe acetate oxidation capacity were treated with ^13^C-labeled acetate (500 mg kg^-1^ body mass) via oral gavage following light anesthesia (4% isoflurane-O_2_ mixture). Acetate oxidation capacity was measured using the CRDS-based breath analysis, after which squirrels were euthanized. Antibiotics were administered as described in [2].

The 11 squirrels used for metagenomic-based estimates of mucin glycan degradation were administered saline via oral gavage following light anesthesia (4% isoflurane-O_2_ mixture). On the day of sampling CRDS-based breath analysis was performed, after which squirrels were euthanized.

All experiments were performed on squirrels whose body temperatures (T_b_) were in the euthermic range (above 30⁰C) so that the physiological and biochemical processes we examined could proceed at optimal temperatures. The T_b_ range of Summer squirrels when euthanized was 36-38 ⁰C. Hibernating squirrels were induced to arouse from a torpor bout (thus simulating the natural interbout arousal periods that last 12-24 h) by transferring the animal cage from the 4°C cold room to the ∼22°C lab. Experiments commenced when T_b_ was at least 36°C.

### Tissue harvesting and preparation for analyses

Following euthanasia, we collected host tissues (liver, quadriceps) and cecal contents (a proxy for the cecal microbiota) and prepared them for isotope ratio mass spectrometry (IRMS) and nuclear magnetic resonance (NMR)-based metabolomics analyses as described in [2]. The IRMS and NMR techniques were executed as described in [2]. All ^1^H and ^13^C spectra were analyzed using the Chenomx software package (Version 8.5; Edmonton, AB, Canada).

### Metagenomics

Generation of metagenomic data from cecal contents was performed as previously described [2]. Briefly, genomic DNA was extracted using the Qiagen DNeasy PowerSoil Kit (Qiagen, Hilden, Germany) according to manufacturer’s specifications and quantified using a Qubit Fluorometer (Invitrogen, San Diego, CA, USA). Sequencing was performed on an Illumina NovaSeq6000 (Illumina, San Diego, CA, USA) using an S4 Reagent Kit v1.5 at 2×150bp PE. Trimmomatic v0.38 [5] was used to remove sequencing adapters and low-quality reads, while host DNA was filtered using bowtie2 v2.2.2 [6] against the 13-lined ground squirrel genome (GenBank and RefSeq assembly accession = GCA_000236235.1). The remaining reads were assembled using metaSPAdes v3.13.0 [7] and open reading frames (ORFs) were predicted using prodigal v2.6.3 [8]

For the carbon fixation-related analyses, translated proteins from the ORF predictions were compared against the Pfam [8] profiles for enzymes involved in Wood-Ljungdahl pathway (WLP) carbon fixation, as defined by KEGG [9–11] using HMMER [12]. These enzymes included fdhF (formate dehydrogenase), fhs1 (formate tetrahydrofolate ligase), fchA (methenyltetrahydrofolate cyclohydrolase), foID (methylenetetrahydrofolate dehydrogenase (NADP+) / methenyltetrahydrofolate cyclohydrolase), metF MTHRF (methylenetetrahydrofolate reductase (NADH)), cdhE acsC (acetyl-CoA decarbonylase/synthase, CODH/ACS complex subunit gamma), cdhD acsD (acetyl-CoA decarbonylase/synthase, CODH/ACS complex subunit delta), cdhC (acetyl-CoA decarbonylase/synthase subunit beta), and cooS acsA (anaerobic carbon-monoxide dehydrogenase catalytic subunit). Identified sequences were then mapped back to their contig of origin and taxonomically classified using centrifuge [13]. The percentage of target genes in the metagenomes were calculated by dividing the number of identified target genes by the total number of predicted genes. The percentage of target genes classified to different taxa was calculated by dividing the number of target genes classified to a specific taxon by the total number of detected target genes.

For the mucin degradation analysis, we first identified putative enzymes thought to be involved in mucin degradation from the Carbohydrate-Active enZymes (CAZy) database [14], including glycoside hydrolase (GH) 20 (exo-acting *β*-*N*-acetylglucosaminidases, *β*-*N*-acetylgalactosamindase and *β*-6-SO_3_-*N*-acetylglucosaminidases), GH33 (sialidase), GH84 (*β*-*N*-acetylglucosaminidases), GH85 (Endo-β-N-acetylglucosaminidases), and GH109 (α-*N*-acetylgalactosaminidase). We then compared the translated proteins from the predicted ORFs of the metagenomes against the CAZy database using dbCAN [15] and HMMER v3.3 [12].

To determine differences in mucin degradation genes between Summer and Late Winter metagenomes, we first normalized CAZyme mucin degradation genes by calculating reads per kb per genome equivalent (RPKG) [16]. For each ORF, ORF length (kb) and the number of mapped short reads were determined using bowtie2 [5]. Next, the number of mapped short reads for each ORF was divided by ORF length and summed in each sample. This sum was then divided by genome equivalents to calculate RPKG. Genome equivalents was determined using MicrobeCensus [16] with default parameters. Seasonal differences in CAZyme normalized counts were analyzed in R [17] using an ANOVA (*aov*) and Tukey’s HSD (*TukeyHSD*) if they were normally distributed, or with the Kruskal-Wallis test (*kruskal.test*) and Dunn’s test (*dunn.test*) if they were not normally distributed. P-values were adjusted for false discovery rate using the Benjamini-Hochberg procedure (*p.adjust*). Results were considered significant if adj P < 0.05, except for Dunn’s test where results were considered significant if adj P ≤ 0.025. Overrepresented CAZymes were identified with DESeq2 [18] and normalized counts were transformed using the Hellinger method (*hellinger*, labdsv package in R).

### Statistical analyses

Multivariate ^13^C-metabolomic data for cecal contents, liver and muscle were analyzed by combining principal components analysis (PCA) for visualization and PERMANOVA analysis for quantitation (MetaboAnalyst 6.0, www.metaboanalyst.ca). Effects of season and microbiota presence on ^13^C-incorporation into identifiable metabolites (via NMR) and protein fraction of tissue (via IRMS) were analyzed using 2-way ANOVA (GraphPad Prism v10.3). Breath δ^13^C was plotted as a function of the 3 h following labeled urea or acetate treatment, and then area-under-the-curve (AUC) was calculated and expressed relative to each individual’s total breath analysis time [19]. The data for urea-treated squirrels originally reported in [2] were expressed here as an exponential function of base *e* to eliminate negative values and enable urea-to-acetate conversion efficiency calculations, which were expressed as the ^13^C-acetate abundance relative to the exponentially converted breath δ^13^C AUC for the urea-treated squirrels. Time course traces for breath responses to doubly labeled urea-treated squirrels can be found in [2]. Seasonal differences in breath δ^13^C AUC for urea-treated squirrels and relative abundances of gut bacterial genes involved in carbon fixation (via WLP) and mucin degradation were analyzed using Student’s t-test. Effects of season and microbiota presence on breath δ^13^C AUC for acetate-treated squirrels were analyzed using 2-way ANOVA. Non-normally distributed data via Shapiro-Wilk test were log-transformed prior to analysis. In cases where data remained non-normal, non-parametric tests were used. For data presented in whisker plots, the upper and lower bounds of the box represent 75^th^ and 25^th^ percentiles, respectively; the horizontal line within the box represents the median value, and the upper and lower whiskers represent the maximum and minimum values within the data set, respectively. Statistical significance was set at P<0.05 for all comparisons.

## Results

### Urea carbon (^13^C) incorporation into host and microbe metabolome and protein

We observed a total of 10, 12 and 15 metabolites enriched in urea-derived ^13^C in cecal contents, liver and muscle, respectively (Figs. 1, 2). The paired PCA-PERMANOVA multivariate analysis revealed that, among the six treatment groups (Summer, Early Winter, Late Winter; with and without microbiotas) within each compartment, ^13^C-enriched metabolite pools varied with season in cecal contents (F=3.0992, R^2^=0.4025, P=0.007) and liver (F=2.7784, R^2^=0.3766, P=0.009), but not in muscle (F=1.2785, R^2^=0.2175, P=0.288) (Fig. S1). However, among all observable metabolites, only cecal content ^13^C-acetate was significantly affected by the presence of a microbiota, whereby its concentrations in microbiota-intact squirrels were 2-to 7-fold higher than those of microbiota-depleted squirrels (Fig. 1A; ANOVA P<0.0001; original data were reported in Fig. S6 in [2] and expressed here in concentration). This microbiota effect confirmed that the incorporation of urea carbon into acetate molecules was dependent upon microbial activity, most likely reflecting reductive acetogenesis that used ^13^CO_2_ generated by ureolytic bacteria after administration of ^13^C,^15^N-urea. ^13^C-acetate levels in liver were low and unaffected by the gut microbiota, though concentrations were slightly higher in Late Winter than Summer or Early Winter (Fig. 1B; ANOVA P>0.05). In muscle, ^13^C-acetate levels were below detectable limits.

**Figure 1.**
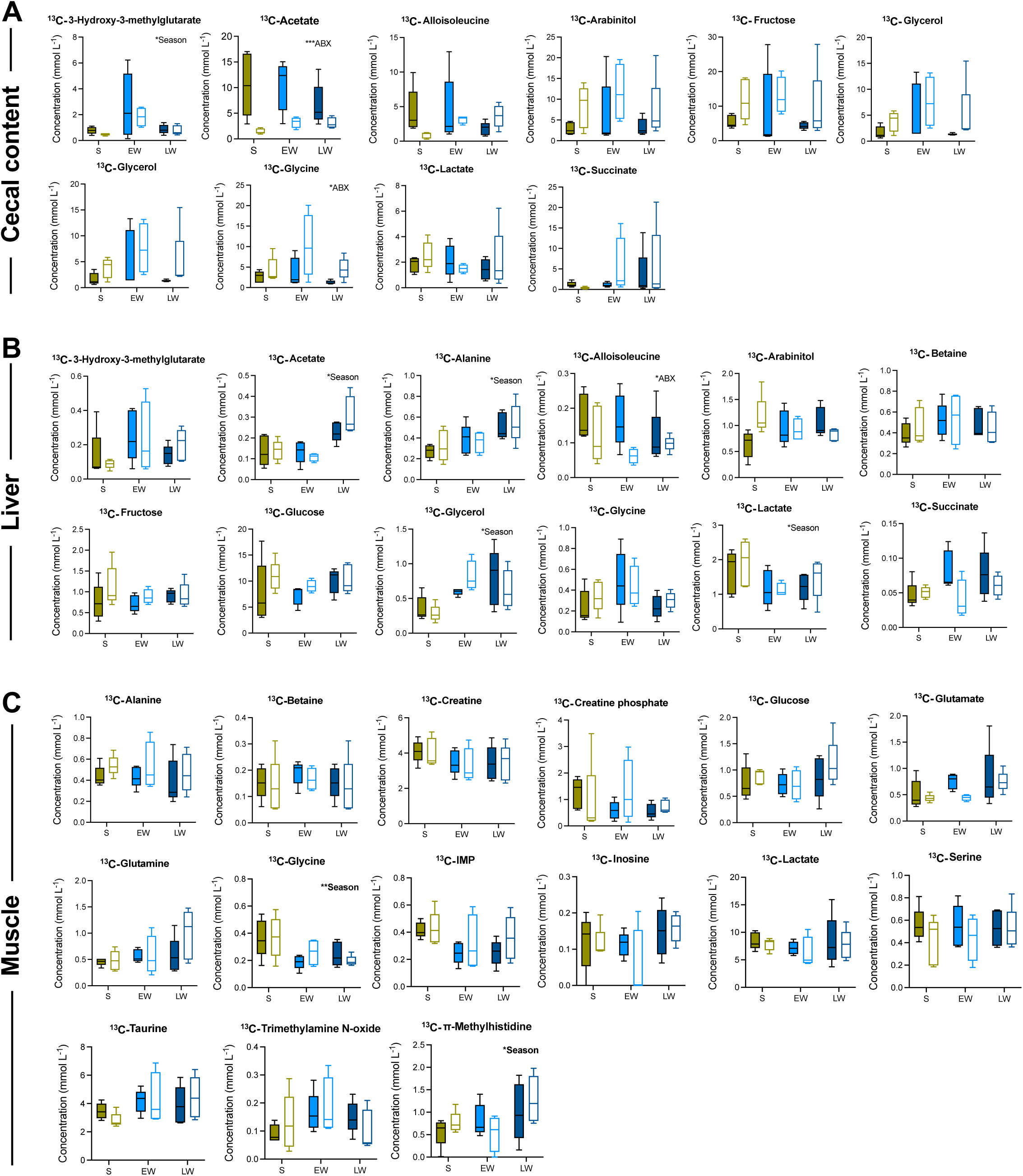
^13^C-urea carbon incorporation into metabolomes for cecal contents (A), liver (B), and quadriceps muscle (C) of microbiota-intact (filled bars) and microbiota-depleted (open bars) 13-lined ground squirrels from Summer (S), Early Winter (EW) and Late Winter (LW). For each panel, the plots portray ^13^C-incorporation into the metabolite named above the plot. Asterisks indicate a significant effect of season (Season) or gut microbiota presence/absence (ABX), where * means P<0.05, ** means P<0.01, and *** means P<0.001. n=5 for all data.

In contrast to our previous observations [2] that urea-derived ^15^N is incorporated into host (liver, muscle) and microbiota (cecal contents) protein pools, we found no evidence of microbiota-dependent incorporation of ^13^C into protein fractions of the liver, muscle, or microbiota after ^13^C,^15^N-urea administration (Fig. S2). For the microbiota-intact squirrels, the small volumes of cecal content samples required pooling all samples into a single aliquot for IRMS analysis, which ultimately precluded statistical testing. For microbiota-depleted squirrels, the samples volumes were too small for analysis. However, we observed a trend for greater incorporation of ^13^C into protein fractions of Summer cecal contents compared with Early or Late Winter fractions (Fig. S2).

### Seasonal effect on ureolytic activity and urea-to-acetate efficiency

The CRDS-based measurements of breath δ^13^C reported in [2] for estimation of in vivo gut microbial ureolytic activity were transformed here to express the δ^13^C values as positive numbers, thus enabling efficiency measurements (see Materials and Methods). This analysis revealed that, as reported in [2], microbial ureolytic activity was higher in Summer than in Early or Late Winter squirrels, which were not different (Fig. 2A; ANOVA P=0.0017).

**Figure 2.**
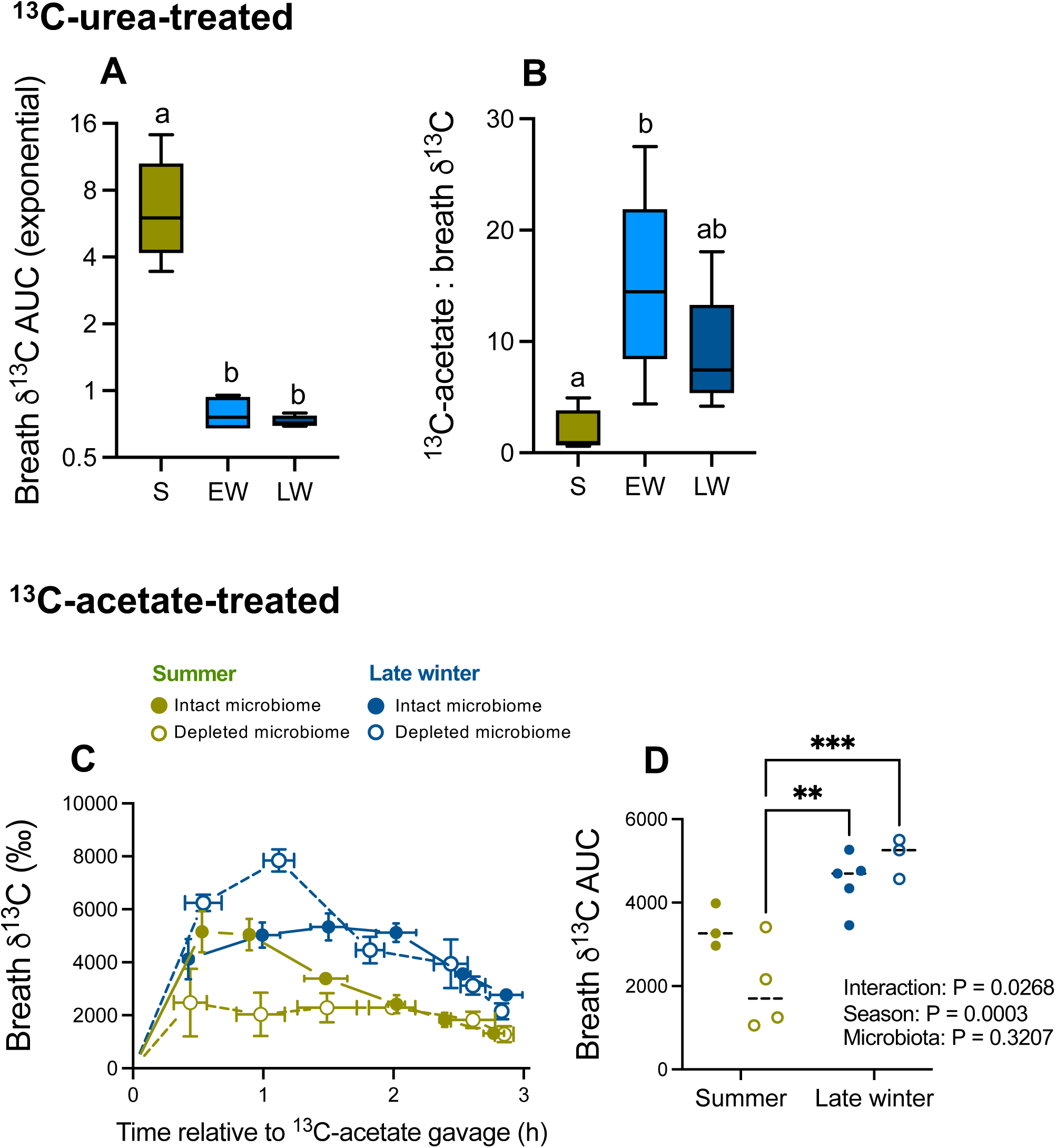
(A) Breath δ^13^C values for 13-lined ground squirrels from Summer (S), Early Winter (EW) and Late Winter (LW) treated with ^13^C-labelled urea. The data represent total area under the curve (AUC) for the 3 h period following urea treatment, as first reported in [2]. The data are expressed here exponentially to eliminate negative values and thus enable the urea-to-acetate efficiency calculations. (B) Urea-to-acetate efficiency, which is the ratio of an individual squirrel’s ^13^C-enriched cecal acetate concentration to its breath δ^13^C AUC value. (C) Mean breath δ^13^C for Summer (green) and Late Winter (blue) 13-lined ground squirrels treated with ^13^C-labeled acetate via oral gavage at time 0. Horizontal and vertical error bars represent SEM; closed and open symbols represent squirrels with intact and depleted microbiotas. (D) Mean area under the curve (AUC) values for data in (C), with 2-way ANOVA results presented on the graph and asterisks indicating significant differences as determined by post-hoc test, where * indicates P<0.05, ** indicates P<0.01, and *** indicates P<0.001. n=5 for A and B, n=3-4 for C and D.

Each squirrel’s urea-to-acetate efficiency was expressed as the ratio of its cecal urea-derived ^13^C-acetate concentration to its total ^13^CO_2_ abundance, represented as its transformed breath δ^13^C value shown in Fig. 2A. This efficiency measure reflects the amount of liberated urea carbon that is directed towards acetogenesis. We found that urea-to-acetate efficiency is generally higher during hibernation than summer (Fig. 2B; ANOVA P=0.0135; t-test P=0.0112 when grouping Early and Late Winter into a single hibernation group).

### Seasonal effect on microbial carbon fixation capacity

To estimate the capacity of the squirrel gut microbiome to fix carbon and enable acetogenesis, we analyzed Summer and Late Winter microbial metagenomes for nine genes encoding for WLP-related enzymes. The relative abundances of eight out of nine genes were sustained from Summer to Late Winter (Fig. 3A) despite the ∼90% reduction in bacterial density by Late Winter [20], while the ninth fchA (methenyltetrahydrofolate cyclohydrolase) increased from Summer to Late Winter (Fig. 3A; t-test P=0.003). Analysis of the top 10 bacterial genera from taxonomic classification of WLP genes, as determined by sequence relative abundance, revealed a seasonal shift from Summer to Late Winter in the bacterial potential for reductive acetogenesis that could include use of urea carbon (Fig. 3B). Most notable was the finding that the greatest percentage of WLP genes in Summer metagenomes (1.74%) belonged to *Bacteroides*, whereas over twice that amount (3.63%) was classified to *Blautia* in Late Winter. The percentage of WLP gene sequences represented in *Blautia* spp. in Late Winter metagenomes was over 5-fold than in Summer metagenomes.

**Figure 3.**
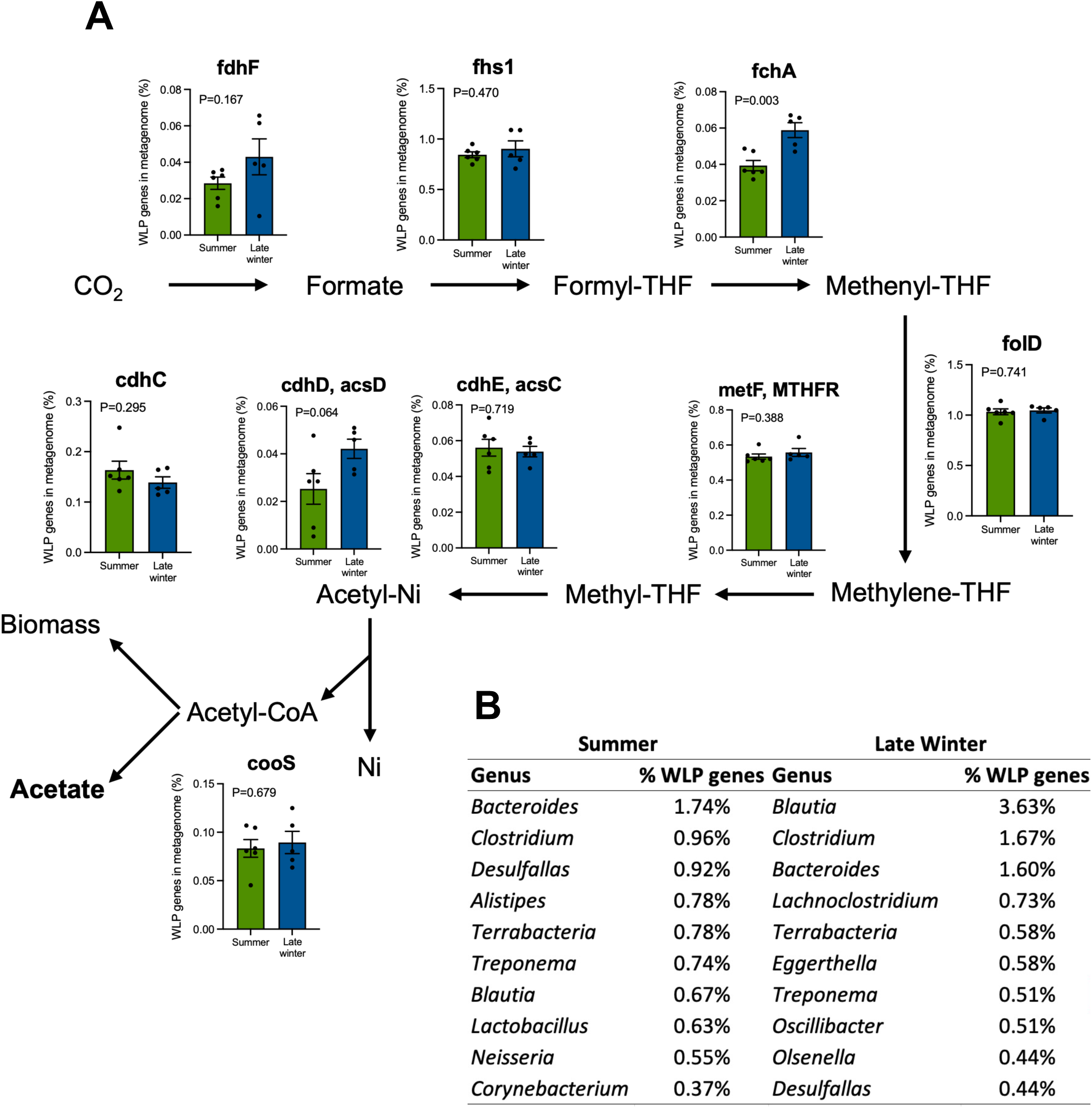
Seasonal effect on (A) the percentage of nine Wood-Ljungdahl pathway (WLP)-related gut bacterial genes for carbon fixation (see text for abbreviation definitions) in the 13-lined ground squirrel gut microbiome, as well as (B) the top 10 bacterial genera from taxonomic classification of WLP genes as determined by sequence abundance.

### Capacities for acetate oxidation

An increase in δ^13^C breath after administration of ^13^C,^15^N-urea indicates that oxidation of one or more bioenergetic metabolites by the microbiota, by squirrel cells, or both is a consequence of ureolysis in the gut lumen. Our identification of ^13^C-acetate molecules in the microbial metabolome that contain urea-derived carbon [2] implicated acetate as one of those metabolites. To explore the site of acetate oxidation in the squirrel holobiont, and whether there is a seasonal difference in capacity for acetate oxidation by the microbiota and/or host, we orally gavaged microbiota-intact and-depleted squirrels from Summer and Late Winter with a body mass-specific dose of ^13^C-labeled acetate and then measured their breath δ^13^C using CRDS. None had been treated with labeled urea. We found that squirrels from all groups exhibited rapid and large increases in breath δ^13^C (Fig. 2C), indicative of acetate oxidation and subsequent release of ^13^CO_2_ in breath. Squirrels in Late Winter generally exhibited greater breath δ^13^C responses than squirrels in Summer (Fig. 2D; ANOVA P=0.0003), but there were seasonal differences in how the presence of the gut microbiota influenced the responses. In Summer, breath δ^13^C as measured by AUC was highest in squirrels with intact microbiotas, whereas in Late Winter, breath δ^13^C was highest in squirrels with depleted microbiotas (Fig. 2D). Consequently, microbiota-depleted squirrels exhibited both the lowest (Summer) and highest (Late Winter) responses, with a >3-fold difference between the two (Fig. 2D; ANOVA P=0.001) suggesting that the host may be especially primed for acetate absorption and oxidation during the winter fast. Season also influenced the time required to reach peak breath δ^13^C, with Summer squirrels reaching peak values Early (∼30 min post-gavage) relative to Late Winter squirrels (∼75 min post-gavage) (Fig. 2C).

### Mucin degrading capacity in the squirrel microbiome

Like urea, mucin glycoproteins are host-derived compounds that provide benefits to gut microbes and their hosts. Mucins are secreted from goblet cells in the gut epithelial layer and are either bound to epithelial cell membranes or secreted directly into the lumen. Mucin glycans are complex carbohydrates whose degradation releases a variety of smaller oligosaccharides that are fermented anaerobically, producing energy-rich metabolites including SCFAs that can be oxidized for ATP generation or serve other roles such as signalling molecules.

Owing to a lack of commercially available ^13^C-labeled mucins that are resistant to breakdown by mammalian enzymes, we used metagenomics to test the hypothesis that the seasonal shifts in gut microbiota composition we previously identified [21,22] are associated with an enhanced capacity to degrade mucins. We focused on gene families that encode CAZymes with the capacity to degrade dietary polysaccharides, mucin glycans, or both, and identified 17 that were overrepresented in either Summer or Late Winter metagenomes. Of the 17, we found five CAZymes that were enriched in Late Winter vs. Summer, all of which are capable of degrading intestinal mucins (Fig. 4).

**Figure 4.**
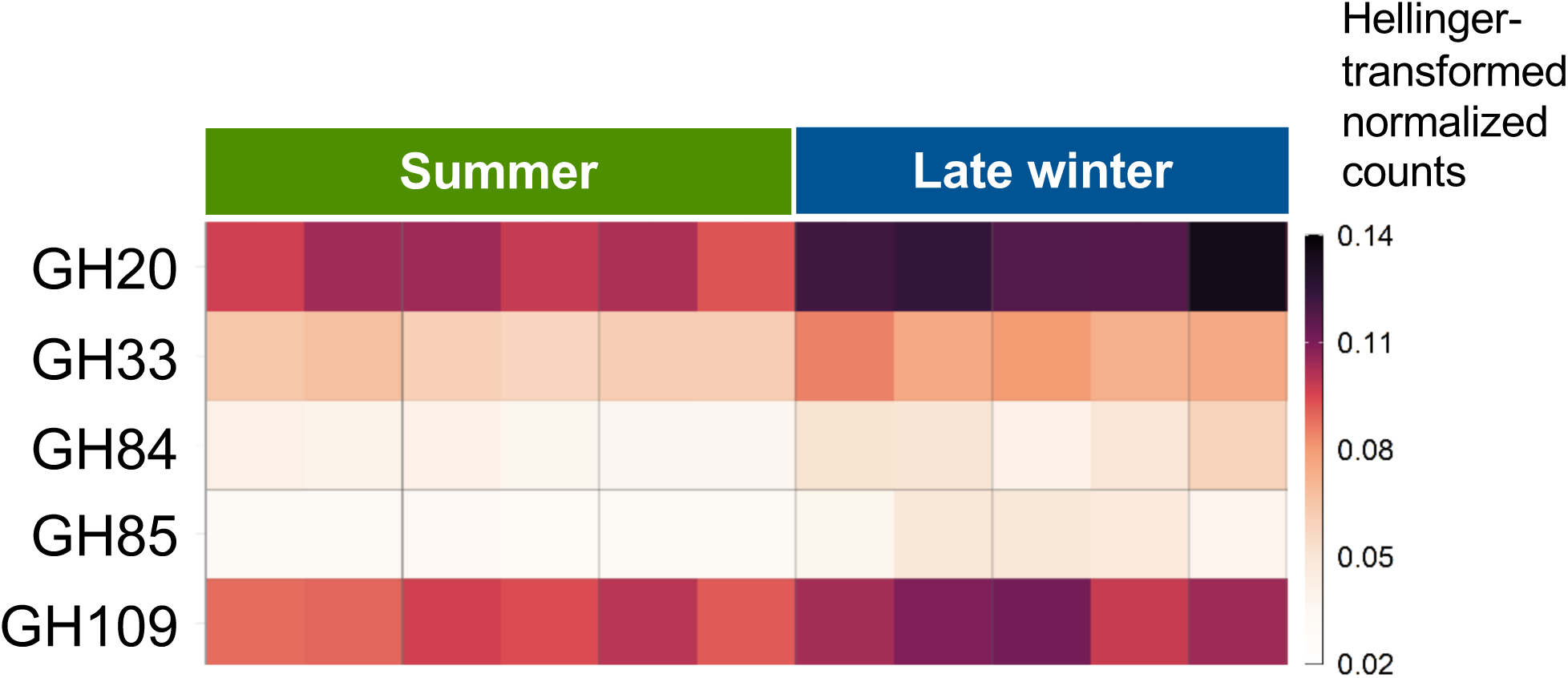
Glycoside hydrolase (GH) carbohydrate-active enzymes (CAZymes) involved in mucin degradation that are seasonally enriched in Late Winter compared with Summer squirrels. Displayed counts represent DESeq2-normalized counts that were Hellinger-transformed. Darker colors indicate higher counts and lighter colors indicate lower counts. T-test results: GH20 P=0.0043; GH33 P<0.0001; GH84 P=0.0048; GH85 P=0.0027; GH109 P=0.0051. N=6 for Summer, N=5 for Late Winter.

## Discussion

This study was designed to provide functional context for our previous observation that in 13-lined ground squirrels, carbon contained in urea molecules that enter the gut lumen is incorporated into acetate within the microbial metabolome [2]. Here we show that this effect is restricted to acetate among a suite of metabolites we examined and is more efficient during winter fasting than the summer fed season. Together, our findings support the idea that through the symbiotic relationship between hibernators and their gut microbes, urea is repurposed as a dual-use endogenous nutrient source during hibernation, with both its carbon and nitrogen salvaged and used for synthesis of important biomolecules. This previously undescribed pathway for acetate production in the ground squirrel gut uses CO_2_ released by ureolytic microbes for hydrogen disposal via reductive acetogenesis, thereby contributing to maintenance of an efficient fermentative environment for the degradation and use of endogenous host compounds, such as mucins, that benefit the hibernator and its microbiota during the long winter fast.

### Urea carbon salvage

Like UNS, UCS begins with the movement of urea molecules from the blood into the gut lumen, mediated by urea transporters (UT-B), where it is hydrolyzed by ureolytic gut microbes. We recently showed that the abundance of UT-B in the squirrel cecum increases from Summer to Late Winter, suggesting a greater capacity to transport urea into the lumen during the hibernation season [2]. We also showed that ureolytic bacteria are found in the squirrel microbiota and that rates of ureolysis are higher in Summer than in Early or Late Winter, likely due to the reduction in total bacterial abundance that occurs in ground squirrels from summer to winter [20,21].

Urea carbon is next incorporated into organic compounds, a process carried out by acetogenic microbes. These metabolites then become available for use by microbes and/or absorption and processing by the host. To investigate this, we screened metabolomes of cecal contents (a proxy for the microbiota) and the squirrels’ liver and skeletal muscle specifically for ^13^C-enriched metabolites which, because our squirrels were treated with ^13^C,^15^N-labeled urea, would indicate urea carbon-containing metabolites. Looking broadly across the multivariate data via paired PCA and PERMANOVA analyses, we found no significant effect of gut microbiota presence/absence on the ^13^C-metabolomes of these three compartments (Fig. S1), suggesting that at this level of analysis there is minimal incorporation of urea carbon into organic compounds. However, analysis of individual metabolites revealed a significant gut microbe-dependent incorporation of urea carbon into the SCFA acetate (Fig. 1).

Our finding that urea carbon-derived acetogenesis occurs in months-long fasted hibernating mammals was notable for several reasons. First, although cecal acetogenesis by the reduction of CO_2_ with H_2_ was first reported in 1977 [23] and replicated subsequently in the hindgut of other monogastric species, the carbon source in those studies was not identified and generally assumed to be oligosaccharides produced by fermentation of dietary carbohydrates. Recently, Firth et al. [24] reported that colonization of mouse and human microbial communities with a ureolytic strain of the gut microbe *Blautia* leads to urea-dependent acetogenesis and cross-feeding of urea carbon from acetate to other SCFAs. Our study demonstrates urea carbon-incorporation into acetate in a wild species, extends its physiological relevance by initiating the experiments with circulating urea, and proposes that the host-microbe interactions that enable urea carbon-derived acetogenesis are regulated in part by the nutritional status of the host animal.

Also notable is the similarity of cecal ^13^C-acetate concentrations throughout the year despite the decrease in bacterial abundance during hibernation [20]. This is consistent with the sustained percentage from Summer to Winter of bacterial genes that encode for enzymes in the Wood-Ljungdahl pathway (WLP; Fig. 3A), though the taxa assigned to those genes differ (Fig. 3B). In this pathway, CO_2_ is reduced by H_2_ to form acetate and water, thus linking the CO_2_ produced by urea hydrolysis with acetogenesis (Fig. 5). Taken together, these findings suggest that the chronic lack of dietary substrates during hibernation induces a compensatory response in the squirrel microbiome that expands metabolic function and enhances survival in those bacteria that persist during winter fasting. Genes encoding carbon fixation-and acetogenesis-related enzymes would facilitate the production of bioenergetic substrates such as acetate that could positively impact both microbes and the hibernator host. This is consistent with the stringent response, whereby bacteria under nutrient stress such as carbon starvation reallocate resources from growth towards the upregulation of survival-related genes until conditions improve [25–27].

**Figure 5.**
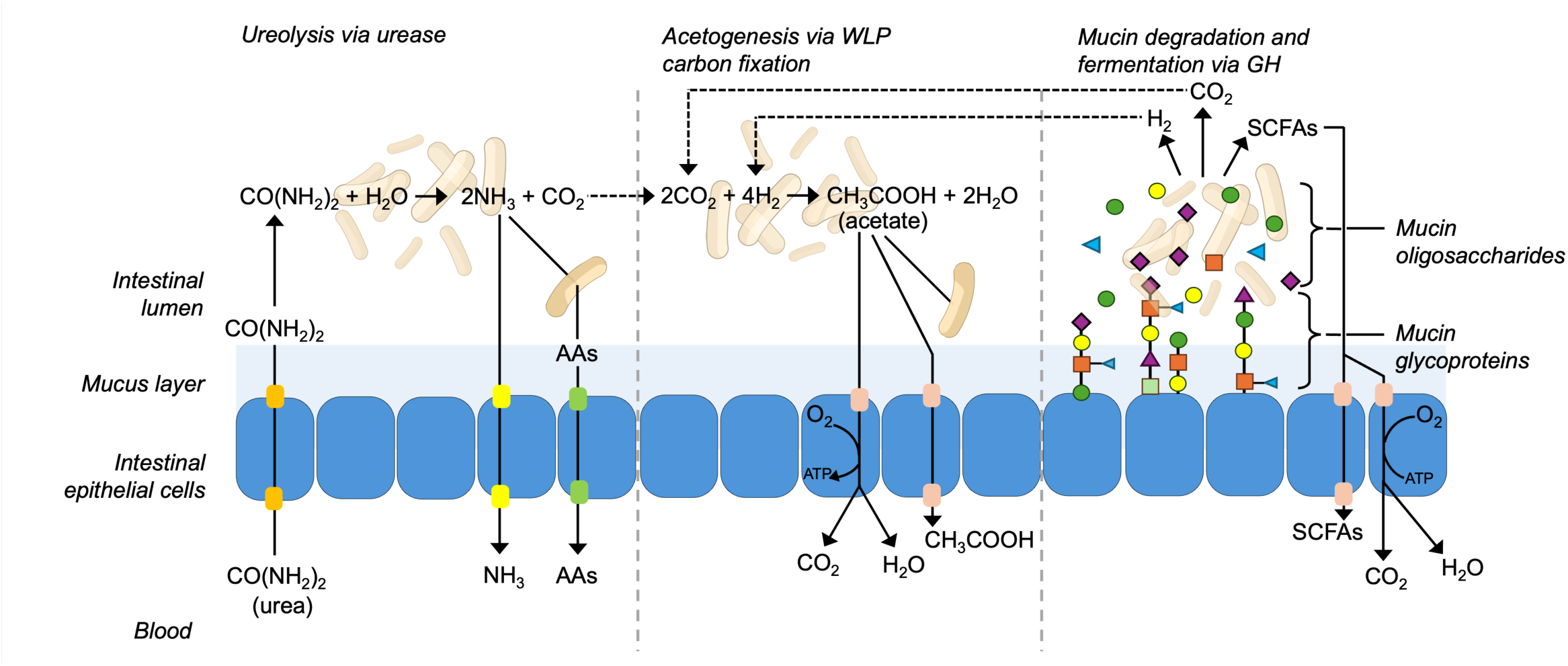
Proposed mechanism for urea carbon salvage in hibernating mammals during euthermic interbout arousals. Urea, endogenously produced in the liver, is transported by epithelial urea transporters from the blood into the cecal lumen, where it is hydrolyzed by ureolytic bacteria into NH_3_ and CO_2_. NH_3_ is subjected to different possible fates, some resulting in urea nitrogen incorporation into microbial and host tissue protein pools via microbial and host amino acid (AA) synthesis, a process called urea nitrogen salvage [2]. CO_2_ serves as the substrate for bacterial Wood-Ljungdahl pathway (WLP) carbon fixation producing acetate, a short chain fatty acid (SCFA) that is oxidized by gut microbes and host cells for energy, as well as serving other potential roles in the host (see text). The WLP reactions consume H_2_, which sustains an efficient fermentation environment for degradation of host compounds such as mucin glycoproteins. Together, this 2-step mutualism transforms urea from a waste molecule into a useful nutrient source and unlocks the value of other endogenous nutrient sources such as mucins for both host and microbe during the resource-limited hibernation season.

Another example of molecular shifts activated by changing nutrient supply is the canonical glucose-acetate transition in *Escherichia coli*, whereby cells excrete acetate while growing on glucose and later absorb and catabolize it for energy when glucose is depleted. This is facilitated by rapid changes in gene expression that suppress biosynthetic pathways and upregulate catabolic ones [28,29]. Here, we posit that a similar strategy may exist for the microbiomes of fasting hibernators; when typical carbon sources in the gut such as glucose derived from dietary polysaccharides are depleted over the winter fast, bacteria scavenge for other carbon sources [28]. Urea carbon-derived acetogenesis appears to be one such pathway that persists throughout hibernation, enabled in part by a sustained (or slightly elevated) proportion of bacterial genes involved in carbon fixation and acetogenesis (Fig. 3A). In the case of fasting hibernators, the contributions of pathways that recycle endogenous host-derived compounds such as urea and mucins may play significant roles in energy-sparing, thus conserving stored lipids that are the major fuel source during torpor and arousals [30].

Interestingly, more of the urea carbon liberated by microbial ureolysis appears to be used for acetogenesis during hibernation than during the summer active season. Specifically, the ratio of ^13^C-acetate to total exhaled ^13^CO_2_ was higher in hibernating than in summer squirrels (Fig. 2B). It is possible that urea carbon is incorporated into compounds other than acetate, but our ^13^C-metabolomics results show that this incorporation is not appreciable, at least in the tissues we analyzed. This suggests that less urea carbon is lost as CO_2_ in breath during winter, when exogenous carbon intake has ceased and when urea carbon, via carbon fixation and subsequent acetogenesis, would most benefit the hibernator holobiont.

### Potential benefits of urea carbon-derived acetate for hibernator host and microbiota

We perceive several benefits of urea carbon-derived acetogenesis for hibernating species during winter fasting. The most direct is acetate’s value as an energy source for host cells, with intestinal epithelial cells (IECs) the most likely to benefit given their dependence on luminal SCFAs such as butyrate to meet energy needs [31]. Under normal fed conditions, butyrate is the preferred energetic substrate for IECs; however, when butyrate availability is limited, acetate oxidation can supply them with ATP [32]. A similar shift in the nutritional landscape occurs during hibernation, when cecal concentrations of all SCFAs decrease relative to Summer levels, though to different extents. For example, cecal butyrate concentrations fall to as low as 3% of Summer levels, whereas acetate concentrations remain at 30-40% of Summer levels [20,21]. It is therefore possible that urea carbon-derived acetogenesis contributes to this energy supply for the host, particularly at the level of IECs.

To lend empirical support for acetate’s role as an energy source for hibernators and their gut microbes, we orally gavaged Summer and Late Winter squirrels with ^13^C-acetate. We then used CRDS to measure breath δ^13^C, whereby elevated values above baseline indicate oxidation of ^13^C-acetate to ^13^CO_2_. Breath δ^13^C rapidly increased following ^13^C-acetate gavage in microbiota-intact Summer and Late Winter squirrels, resulting in similar AUC values for the 3 h post-gavage collection period (Fig. 2D) despite the lower total bacterial abundance of Late Winter relative to Summer squirrels [20]. To distinguish host acetate oxidation from potential microbial acetate oxidation, we repeated the experiments on microbiota-depleted squirrels in both seasons. In Summer squirrels, depleting the microbiota halved the breath δ^13^C AUC (suggesting a strong microbial contribution to acetate oxidation); whereas in Late Winter squirrels, microbiota depletion had no effect on breath δ^13^C AUC but produced a higher δ^13^C peak that was attained more quickly. Overall, the Late Winter microbiota-depleted squirrels had AUC values 3-fold higher than Summer microbiota-depleted squirrels (Fig. 2D). These results suggest two things about acetate use in hibernating squirrels. First, during summer, gut microbes oxidize much of the acetate they produce (perhaps contributing to greater ^13^C incorporation into cecal content protein; Fig. S2), whereas during hibernation, a greater proportion of the acetate is absorbed and oxidized by the host. Thus, the squirrel host appears to benefit most from the bioenergetic contributions of acetate during hibernation. We suspect that the primary site of oxidation for microbially generated acetate in squirrels, especially during hibernation, is within IECs. This is consistent with the lack of observable ^13^C-acetate in liver and muscle, and with the generally deficient energy state of IECs during long periods of food deprivation [33]. Second, the 3-fold higher breath δ^13^C responses after ^13^C-acetate gavage in Late Winter compared with Summer microbiota-depleted squirrels indicate that the capacity to absorb and oxidize acetate increases during hibernation, when resources are limited and organic compounds such as acetate are especially valuable. Although we have not directly measured rates of intestinal acetate absorption, these results are consistent with our previous work showing higher normalized rates of Na^+^, glucose, and amino acid absorption by the small intestinal epithelium in hibernating than in summer squirrels [34,35]. Similar results were observed for nutrient uptake into brush border membrane vesicles [36], and it is plausible that similar seasonal effects on intestinal acetate absorption via monocarboxylate transporters underlie the results seen here. Another indirect way that urea carbon-derived acetate may influence IEC metabolism during winter fasting is through its conversion to their preferred fuel source by acetate-consuming, butyrate-producing bacteria within the microbiota [37]. Together, these results suggest that the hibernator’s physiology is seasonally optimized to benefit from urea carbon-derived acetogenesis, which may suggest positive selection on this mechanism.

Another potential role for urea carbon-derived acetate in the hibernating host is its role as a ketogenic substrate. The ketone β-hydroxybutyrate is the preferred fuel source for the heart and brain of ground squirrels during hibernation [38]. Ketogenesis, which takes place primarily in the liver but can occur in other sites including the gut [39,40], is regulated by the rate-limiting enzyme (3-hydroxy-3-methylglutaryl-CoA synthetase 2; HMGCS2). We showed previously that HMGCS2 is expressed in the 13-lined ground squirrel intestine and its protein abundance is 2-fold higher during hibernation than in summer [41]. This finding is consistent with those from mice and rats whereby short-term fasting elevates both intestinal gene expression of HMGCS2 [42] and the conversion of acetate to ketone bodies in isolated colonocytes [32]. Furthermore, a role for the microbiota in stimulating hepatic ketogenesis in mice under nutrient deprivation was shown when mice with intact gut microbiotas elevated hepatic ketogenesis following a 24 h fast, but the response was reduced in mice lacking a microbiota [43]. Given that levels of fatty acids, which are the most common substrates for ketogenesis, are likely reduced in hibernator IECs due to the lack of dietary lipid intake, acetate may be an effective ketogenic precursor for the gut during hibernation.

In addition to providing acetate as an energy source, acetogenesis also contributes to maintenance of a gut environment that is conducive to microbial breakdown of dietary substrates and host-secreted compounds. Pathways associated with microbial fermentation require electron sinks to dispose of free elections, and production of H_2_ is a major electron sink. However, high H_2_ partial pressure in the lumen can be harmful to bacterial and host cells and can slow key microbial reactions that favor substrate fermentation and the production of SCFA, such as the regeneration of the coenzyme NAD^+^ from NADH [44]. Microbes whose enzymatic repertoire include H_2_ consumption – such as acetogens and methanogens – thus play essential roles in maintaining the gut lumen in a condition permissive to ongoing fermentation of substrates whilst minimizing damage to microbes and their hosts. A second possible role for UCS could thus be to sustain the fermentation of endogenous host compounds. The most abundant of these are secreted mucins, especially during hibernation when IEC proliferation and sloughing are suppressed [45]. Although methanogenesis plays a similar role (but without formation of a bioenergetic compound like acetate), there is no evidence for hydrogenotrophic methanogens (Archaea) in the microbiotas of hibernating mammals including ground squirrels [20–22] and bats [46]. Thus, acetogenesis is likely the primary H_2_ disposal mechanism used in the hindgut of mammalian hibernators.

### Seasonal changes in mucin degradation capacity in the ground squirrel microbiota

Mucins are complex glycoproteins secreted from goblet cells in the gut epithelium that, upon hydration in the lumen, form mucus, a complex viscoelastic matrix that protects the epithelium and serves as a habitat for a diverse assemblage of microbes [47]. Mucins also provide nutritional benefits to gut microbes and their hosts [47], which would be particularly beneficial during winter fasting. In the absence of dietary (typically plant) glycans, host-derived mucins become major substrates that support microbial growth and proliferation, and they contribute to host energetic needs by producing SCFAs that fuel IECs [48]. Coupled with our previous observation that cecal mucin (MUC2) expression continues during hibernation [22], these molecules, like urea, can serve as valuable sources of carbon, nitrogen and energy during the winter fast when exogenous nutrients are absent. Intact mucins are not digestible by mammalian enzymes, but bacteria that express certain glycosyl hydrolases (GHs) can cleave and metabolize glycan linkages, releasing oligosaccharides that are further utilized by the same or other bacteria for energy, and in so doing produce beneficial metabolites for the host [49].

Our metagenomics analysis revealed a strong seasonal influence on the capacity of the ground squirrel microbiome to degrade mucins during winter fasting. The five CAZyme gene families that are enriched in Late Winter compared with Summer microbiomes all have mucin degradation capacity, whereas those enriched in Summer microbiomes include a mix of mucin and non-mucin degraders. This is consistent with our previous findings that several of the bacteria whose relative abundances increase in Winter are known mucin degraders [22,34].

Moreover, all five genes enriched in the Late Winter microbiomes were classified to at least one member of the Bacteroidota; this phylum, whose relative abundance in hibernating squirrels increases over 2-fold from Summer to Winter [21,22], includes the highest number of species with mucin-degrading capacity in mammals. Other noteworthy genera to which winter-enriched gene families are classified include *Akkermansia, Alistipes* and *Blautia*. *Akkermansia muciniphila*, which is the sole member of the phylum Verrucomicrobiota present in the ground squirrel microbiota [20,21], can subsist on mucin alone as its sole carbon and nitrogen source [50]. This property may be responsible for the over 4-fold increase in relative abundance of Verrucomicrobiota during hibernation [21,22]. The genus *Alistipes*, a member of Bacteroidota and another prominent mucin degrader [49], also increases during hibernation [21,22]. Beyond its role as a mucin degrader, *Alistipes* has the highest percentage of urease genes in the ground squirrel microbiome during hibernation, increasing nearly two-fold in abundance from summer to late winter [2]. *Blautia*, which has the highest percentage of WLP genes in the Late Winter metagenome (Fig. 4B), is also a mucin degrader [49] and contains four of the five mucin-degrading GHs that we found to be elevated in Late Winter vs. Summer microbiomes (Fig 4).

Moreover, like *Alistipes*, *Blautia* also contains genes that encode for urease in ground squirrels [2]. Taken together with our previous findings [2], the results presented here suggest that several bacterial genera play multiple roles in the processing of the endogenous nutrients urea and mucin that yield potential benefits to the hibernator host and its microbiota in the absence of food intake.

## Conclusion

Collectively, these findings introduce UCS as a mutually beneficial mechanism through which hibernating mammals and their gut microbes meet metabolic demands during prolonged fasting. We propose a process whereby carbon limitation in the gut of fasted hibernators is ameliorated by a sequential set of host-microbe interactions involving ureolytic and acetogenic bacteria (Fig. 5). This two-step mutualism – which could be carried out by the same microbe (e.g., *Blautia* [24]) or by different ones (e.g., *Blautia* and *Alistipes*) – links urea hydrolysis with generation of acetate, providing an oxidizable substrate for gut microbes and the hibernator host. And via H_2_ disposal, it helps sustain the gut ecosystem as an efficient environment for microbial fermentation of host compounds such as mucin glycoproteins, thus unlocking the value of endogenous nutrient sources for both the microbiota and its host. In conjunction with UNS — which facilitates sustained protein synthesis in host tissues during hibernation [2] — UCS establishes urea as a valuable endogenous nutrient source throughout the winter fast and underscores the integral role that gut microbes play in hibernation’s metabolic phenotype.

Consistent shifts in gut microbial composition observed across various hibernating mammals [51,52] and the finding that UCS also takes place in summer squirrels suggest that the dual role for urea in enabling its nitrogen and carbon to be recycled into molecules with nutritional value may be widespread among hibernating and non-hibernating mammals, including humans [24]. Taken together, these insights deepen our understanding of the host–microbiome interactions that underlie metabolic resilience and further highlight the vital roles that the gut microbiota plays for animals enduring extended periods without external nutrient sources.

## 5. ACKNOWLEDGEMENTS AND FUNDING SOURCES

We thank K. Krautkramer and S. Martin for insightful comments on an earlier version of this manuscript. The work was supported by National Science Foundation (NSF) Grant IOS-1558044 to H.V.C., F.M.A.-P. and G.S.; National Institutes of Health (NIH) Grants P41GM136463, P41GM103399 and P41RR002301 to the National Magnetic Resonance Facility at Madison; E.C. was supported by a National Institute of General Medical Sciences of the NIH traineeship T32GM008349 and NSF Graduate Research Fellowship DGE-1747503; M.D.R. was supported by a Natural Sciences and Engineering Research Council (NSERC) Canada Postdoctoral Fellowship to M.D.R. and is currently supported by an NSERC Discovery grant and a Canadian Space Agency Research Model grant.

## Supplemental figures

**Figure S1.**
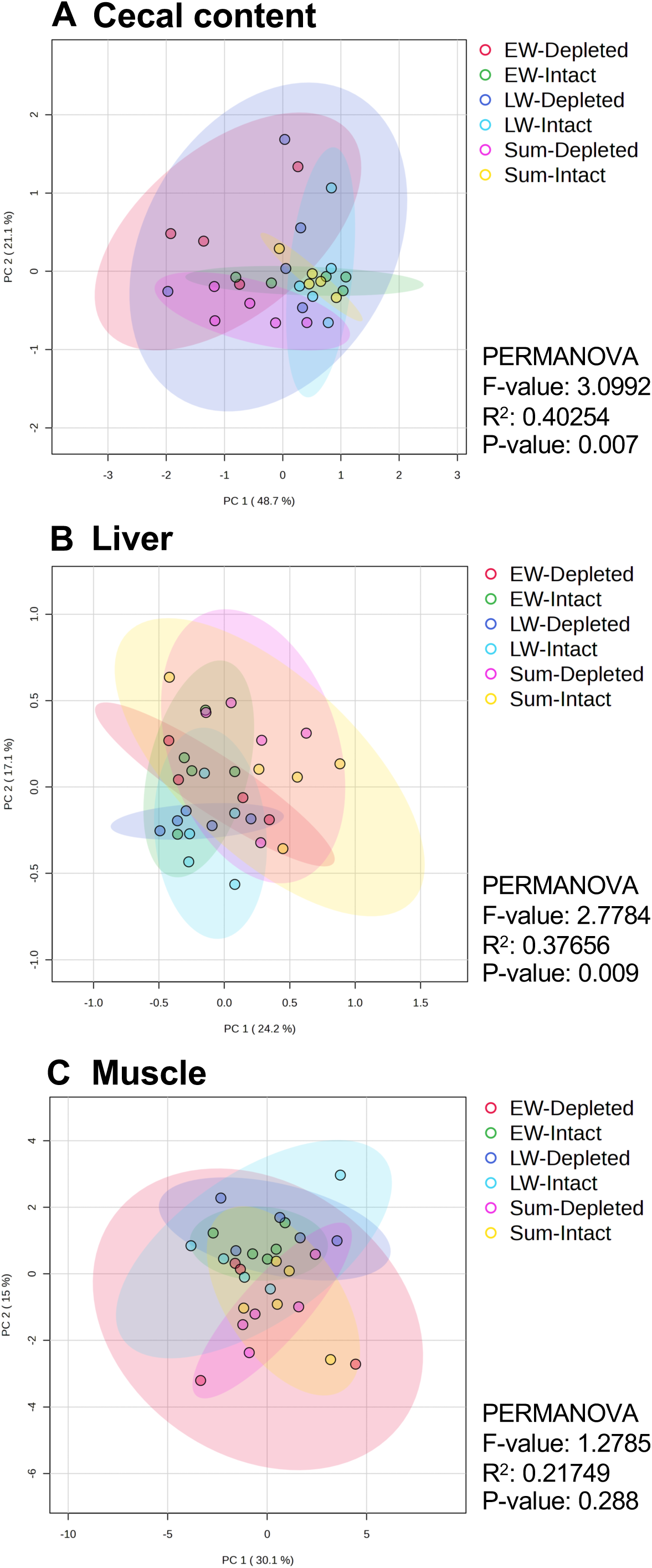
Principal components analyses (PCA) for ^13^C-enriched metabolomes from 13-lined ground squirrel cecal contents (A), liver (B), and quadriceps muscle (C). Treatment groups include gut microbiota-intact and - depleted groups from Summer (sum), Early Winter (EW), and Late Winter (LW) squirrels. PERMANOVA results for each tissue are presented next to its respective score plot.

**Figure S2.**
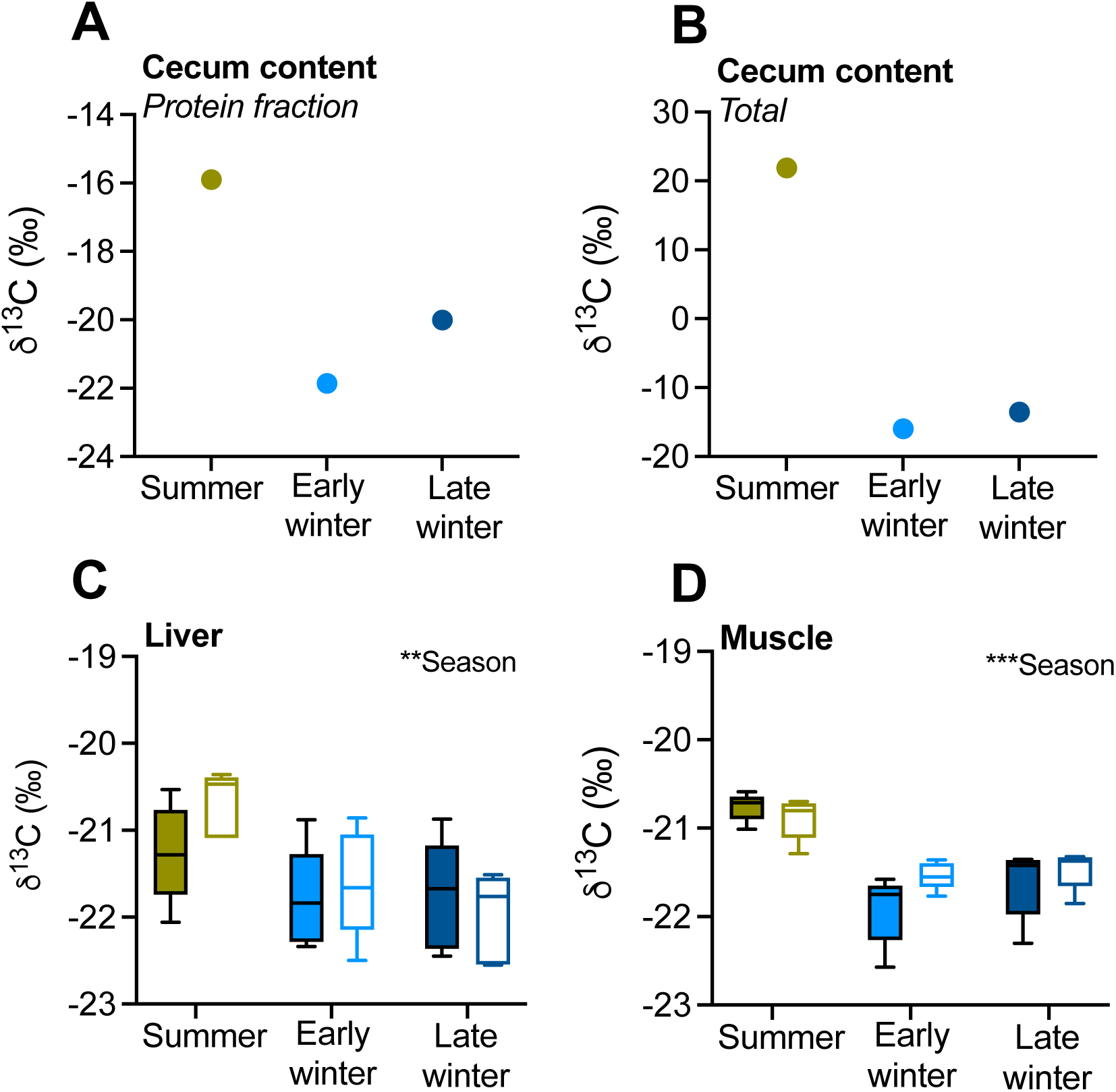
Isotope ratio mass spectrometry-based measurements of ^13^C-urea carbon incorporation into the protein fraction (A) and total milieu (B) of cecum content, as well as protein fractions of (C) liver and (D) quadriceps muscle. Solid bars represent samples from microbiota-intact squirrels, white bars from microbiota-depleted squirrels. Cecum content panels (A, B) lack microbiota-depleted data because ABX-treatment caused insufficient content for analyses. Asterisks indicate a significant effect of season (Season) or gut microbiota presence/absence (ABX), where ** indicates P<0.01 and *** indicates P<0.001. n=5 for all data except for cecum content (A, B; n=1, pooled samples).

## Notes

### Competing Interest Statement

FMA-P is the founder of Isomark Health, Inc.

### Summary of Updates

Figures have been modified from previous versions for style; new summary figure visualizing the proposed multistep mutualism; content added to Discussion; significance statement added.

